# Protected area effectiveness for fish spawning habitat in relation to earthquake-induced landscape change

**DOI:** 10.1101/283333

**Authors:** Shane Orchard, Michael J. H. Hickford

## Abstract

We studied the effectiveness of conservation planning methods for *Galaxias maculatus*, a riparian spawning fish, following earthquake-induced habitat shift in the Canterbury region of New Zealand. Mapping and GIS overlay techniques were used to evaluate three protection mechanisms in operative or proposed plans in two study catchments over two years. Method 1 utilised a network of small protected areas around known spawning sites. It was the least resilient to change with only 3.9% of post-quake habitat remaining protected in the worst performing scenario. Method 2, based on mapped reaches of potential habitat, remained effective in one catchment (98%) but not in the other (52.5%). Method 3, based on a habitat model, achieved near 100% protection in both catchments but used planning areas far larger than the area of habitat actually used. This example illustrates resilience considerations for protected area design. Redundancy can help maintain effectiveness in face of dynamics and may be a pragmatic choice if planning area boundaries lack in-built adaptive capacity or require lengthy processes for amendment. However, an adaptive planning area coupled with monitoring offers high effectiveness from a smaller protected area. Incorporating elements of both strategies provides a promising conceptual basis for adaptation to major perturbations or responding to slow change.

## 1. Introduction

For many species, critical life history phases create obligate habitat requirements. These may be vulnerable points in the life cycle, especially where relatively specific biophysical conditions are required (Lucas, Bubb, Jang, Ha, & Masters, 2009). Vulnerability may be associated with periodic events and longer term change involving both natural and anthropogenic processes (Turner et al., 2003). A particular concern is where human activities reduce the quality or availability of existing habitat unless counterbalanced by compensatory actions, such as the creation of suitable habitat elsewhere (Faith & Walker, 2002). The concept of resilience provides a focus on thresholds in system properties that are important to their persistence (Holling, 1973). In linked socio-ecological systems it is related to adaptive capacity (Gallopín, 2006), and actual responses to changed hazard exposure and/or sensitivity (Turner, Lambin, & Reenberg, 2007). Since resilience assessment is concerned with identifying the conditions required to maintain a desirable state (Gunderson, Allen, & Holling, 2010), it may be readily applied to habitat management.

Protected areas (PAs) describe a desired state defined by clear objectives. They are a cornerstone of global efforts to halt biodiversity loss (UN (United Nations), 2011). The IUCN recognises six categories of PAs defined by differences in management approaches (Stolton, Shadie, & Dudley, 2013). Category IV PAs aim to protect particular species or habitats (Table 1). They are often relatively small and are designed to protect or restore: 1) flora species of international, national or local importance; 2) fauna species of international, national or local importance including resident or migratory fauna; and/or 3) habitats (Dudley, 2008).

**Table 1.** Aspects of IUCN Category IV Protected Areas (Dudley, 2008).

| Role in the landscape/seascape |
| --- |
| Category IV protected areas frequently play a role in “plugging the gaps” in conservation strategies by protecting key species or habitats in ecosystems.<br>They could, for instance, be used to:<br>• Protect critically endangered populations of species that need particular management interventions to ensure their continued survival;<br>• Protect rare or threatened habitats including fragments of habitats;<br>• Secure stepping-stones (places for migratory species to feed and rest) or breeding sites;<br>• Provide flexible management strategies and options in buffer zones around, or connectivity conservation corridors between, more strictly protected areas that are more acceptable to local communities and other stakeholders. |
| Issues for consideration |
| • Many category IV protected areas exist in crowded landscapes and seascapes, where human pressure is comparatively greater, both in terms of potential illegal use and visitor pressure.<br>• The category IV protected areas that rely on regular management intervention need appropriate resources from the management authority and can be relatively expensive to maintain unless management is undertaken voluntarily by local communities or other actors.<br>• Because they usually protect part of an ecosystem, successful long-term management of category IV protected areas necessitates careful monitoring and an even greater than-usual emphasis on overall ecosystem approaches and compatible management in other parts of the landscape or seascape. |

Effective conservation involves managing risks and recent biodiversity declines appear to be continuing (Butchart et al., 2010). Management effectiveness evaluation is an essential activity to assess the strengths and weaknesses of protection mechanisms and different management approaches (Stolton et al., 2007). A key area of focus is the extent to which PAs actually deliver on their objectives such as by protecting important values (Hockings, 2003). Under conditions of environmental change evaluation is especially important to address whether the areas involved are functioning as an effective conservation strategy (Leverington, Costa, Pavese, Lisle, & Hockings, 2010). Various methodologies have been used, many of which were originally developed to the support adaptive management of PA sites and systems (Coad et al., 2015). Range shifts are a topic of particular importance since they may undermine the effectiveness of PA networks unless resilience has been incorporated by design. In this setting human agency is inextricably linked to the trajectory of the values identified for protection. This may require amendment of the protection mechanism itself to ensure continued performance over time.

Diadromous fishes have specific habitat requirements across several stages of their life histories, involving both freshwater and marine environments (Gross, Coleman, & McDowall, 1988). In some species these may be separated by vast distances and associated with significant migrations (Metcalfe, Arnold, & McDowall, 2002). There may be different conservation issues affecting each critical habitat requiring a wide range of management responses (McDowall, 1999). *Galaxias maculatus* (Jenyns 1842) or ‘īnanga’ is a diadromous species currently listed as ‘at risk – declining’ under the New Zealand Threat Classification System (Goodman et al., 2014). Adult fish are found in lowland coastal waterways with the upstream distribution limited by relatively poor climbing ability (Baker & Boubee, 2006; Doehring, Young, & McIntosh, 2012). Spawning occurs in estuarine waterways with the exception of some populations that have become land-locked in lakes (Chapman, Morgan, Beatty, & Gill, 2006). The locations used are highly specific as the result of specialised reproductive behaviour associated with the migration of adult fish towards rivermouths at certain times of the year (Benzie, 1968a). Spawning events are strongly synchronised with the spring high tide cycle with an apparent association between spawning site distribution and the salinity regime (Burnet, 1965). The majority of spawning sites have been found within 500 m of the inland limit of salt water (Richardson & Taylor, 2002; Taylor, 2002). In addition, spawning sites occupy only a narrow elevation range located on waterway margins just below the spring tide high-water mark (Taylor, 2002). As tidal heights drop towards the neap tides these sites are no longer inundated at high-water and for most of their development period the eggs are in a terrestrial environment (Benzie, 1968a, 1968b). Egg survival rates are highly dependent on the condition of the riparian vegetation in these locations until hatching in response to high water levels, usually provided by the following spring tide (Hickford, Cagnon, & Schiel, 2010; Hickford & Schiel, 2011).

The degradation of spawning habitat has been identified as a leading factor in the species’ decline (McDowall, 1992; McDowall & Charteris, 2006). This has been linked to land-use intensification on coastal waterway margins (Hickford et al., 2010), as is a common trend worldwide (Kennish, 2002). Protection mechanisms must often address contested-space contexts characterised by incompatible activities. Multiple-stressor situations are common with grazing, vegetation clearance, mowing, grazing, flood protection, and channelization being examples that have contributed to degradation (Hickford & Schiel, 2011; Mitchell & Eldon, 1991). Habitat protection is a requirement of national legislation under the Conservation Act 1987 and the Resource Management Act 1991 (RMA). Implementation relies on the identification of areas for protection coupled with relevant rules and documented in plans or management strategies prepared under the relevant Acts. In many cases spatial explicit planning methods (e.g. maps) are used to delineate the protected areas. Although these provide a practical approach to address the conservation objective, they require reliable habitat information. In dynamic environments challenges include recognising spatiotemporal variance and accommodating it in design of the protection mechanisms used (Bengtsson et al., 2003).

In 2010 and 2011 a sequence of major earthquakes affected the Canterbury region of New Zealand. It included several large destructive events and numerous aftershocks centred beneath the city of Christchurch (Beavan, Motagh, Fielding, Donnelly, & Collett, 2012). The magnitude of physical effects necessitated a long-term socio-ecological response associated with new ecological trajectories and variety of land-use planning needs. Topographic and bathymetry measurements identified enduring changes in ground levels, especially in the vicinity of waterways (Quigley et al., 2016). Ecohydrological effects have been a particular focus in light of changed water levels on the landscape (Hughes et al., 2015), and alterations to estuarine dynamics (Measures et al., 2011; Orchard & Measures, 2016). *G. maculatus* spawning was recorded at locations never previously utilised in comparison to pre-quake records (Orchard & Hickford, 2016). Vulnerability assessments identified anthropogenic threats at many of these locations and recommended review of protection methods in the operative statutory plans (Orchard, Hickford, & Schiel, in press). This context presented a unique opportunity to evaluate conservation planning options in light of landscape-scale change whilst informing the practical needs of post-quake adaptation processes. The objectives of this paper are to (1) evaluate the efficiency and effectiveness of contemporary protection mechanisms, and (2) identify recommendations for conservation planning to address earthquake-induced landscape change.

## 2. Methods

### 2.1 Study area

The study area is the Avon Heathcote Estuary (Ihutai) located at 43.5°S, 172.7°E in the city of Christchurch (Figure 1). The estuary is located between the Waimakariri River and the southern end of a large sandy bay (Pegasus Bay) where it is a prominent local feature (Kirk, 1979). It is a barrier enclosed tidal lagoon type estuary (Hume, Snelder, Weatherhead, & Liefting, 2007) with high ecological and social values including cultural significance for Māori (Jolly & Ngā Papatipu Rūnanga Working Group, 2013; Lang et al., 2012).

**Figure 1.**
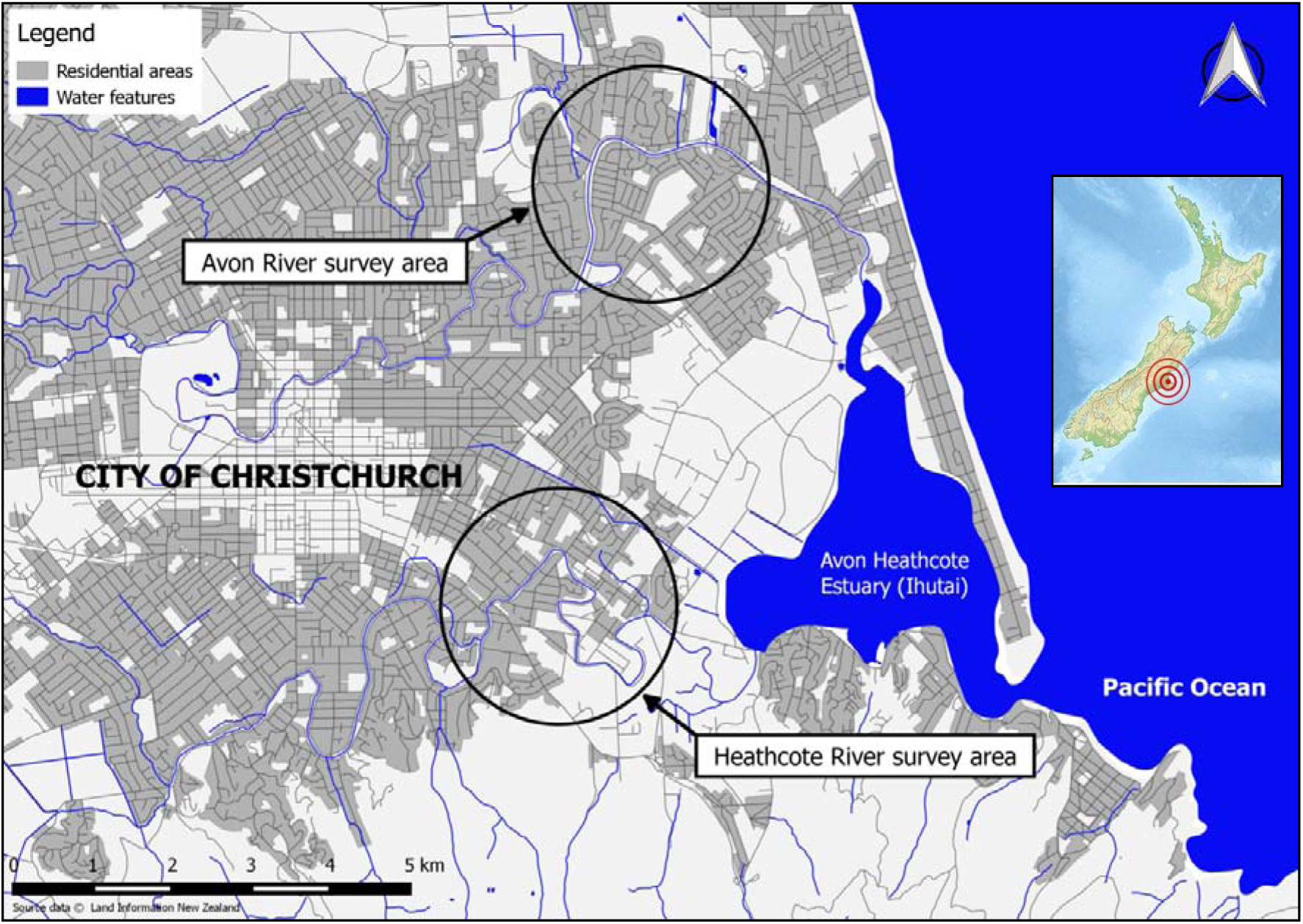
Location of post-earthquake survey areas for *G. maculatus* spawning habitat in the Avon and Heathcote River catchments, city of Christchurch, New Zealand.

The Avon and Heathcote are the two major rivers of the estuarine system, both of which provide *G. maculatus* spawning habitat. These are spring-fed lowland rivers waterways with average base flows of approx. 2 and 1 cumecs respectively (White, Goodrich, Cave, & Minni, 2007). They are also among the most well studied spawning locations in New Zealand with surveys having been conducted periodically since 1988 (Taylor, Buckland, & Kelly, 1992).

### 2.2 Geospatial analyses

We analysed spawning site data from post-earthquake studies comprising of seven independent surveys conducted over two years during the peak spawning months using a census-survey methodology designed to detect all spawning in the catchment (Orchard & Hickford, 2017). The areas surveyed were approximately 4 km reaches in each river extending from the saltmarsh vegetation zone (downstream), to 500 m upstream of the inland limit of saltwater (Figure 1). The dataset of 188 records provided details of 121 spawning occurrences in the Avon and 67 in the Heathcote. Each record included upstream and downstream coordinates of the spawning site, mean width of the egg band, and area of occupancy (AOO) of eggs, with each site being defined as a continuous or semi-continuous patch of eggs (Orchard, Hickford, & Schiel, 2016; Orchard et al., in press).

Three spatially explicit protection mechanisms were identified in an analysis of proposed and operative resource management plans (Table 2). In this paper we use the term ‘protected areas’ to denote spatially explicit areas identified in planning methods to address conservation objectives in statutory policies and plans. The areas evaluated in this study are consistent with the IUCN definition of Category IV protected areas being ‘areas to protect particular species or habitats, where management reflects this priority’ (Dudley, 2008). The size of these areas is often relatively small with varying management arrangements depending on protection needs (Stolton et al., 2013).

**Table 2.** Protected area mechanisms for *G. maculatus* spawning habitat evaluated in this study.

| Method | Protected area mechanism | Delineation method in plans | Information source | Planning documents |
| --- | --- | --- | --- | --- |
| 1 | Network of small protected areas based on known spawning sites | 20 m diameter areas centred on point data coordinates of known spawning sites, identified in schedule to the plan | Point data and descriptions from NISD<sup>†</sup> and historical reports (Maw & McCallum-Clark, 2015) | Environment Canterbury (2015)<br>Environment Canterbury (2016) |
| 2 | Mapped reaches of potential spawning habitat on a catchment basis | Reaches identified in planning maps and referenced in the plan | NISD point data and historical reports coupled with field surveys of riparian vegetation to identify potential habitat (Margetts, 2016) | Environment Canterbury (2014) |
| 3 | Mapped polygons of predicted spawning habitat coupled with a text description of where in the polygon the protection requirements apply | Polygons identified in planning maps and GIS layer referenced in the plan | GIS based model of predicted spawning habitat (Greer, Gray, Duff, & Sykes, 2015) | Environment Canterbury (2017) |

<sup>†</sup> National Inanga Spawning Database

Protected area and spawning site data were visualised in QGIS v2.8.18 (QGIS Development Team, 2016) and reach lengths (RL) calculated in relation to the centrelines of waterway channels digitised from 0.075 m resolution post-quake aerial photographs (Land Information New Zealand, 2016). Three comparable RL metrics were calculated to reflect (a) the RL protected under each planning method, (b) extent of occurrence (EOO) of spawning sites, and (c) the total AOO of spawning sites (Table 3).

**Table 3.**
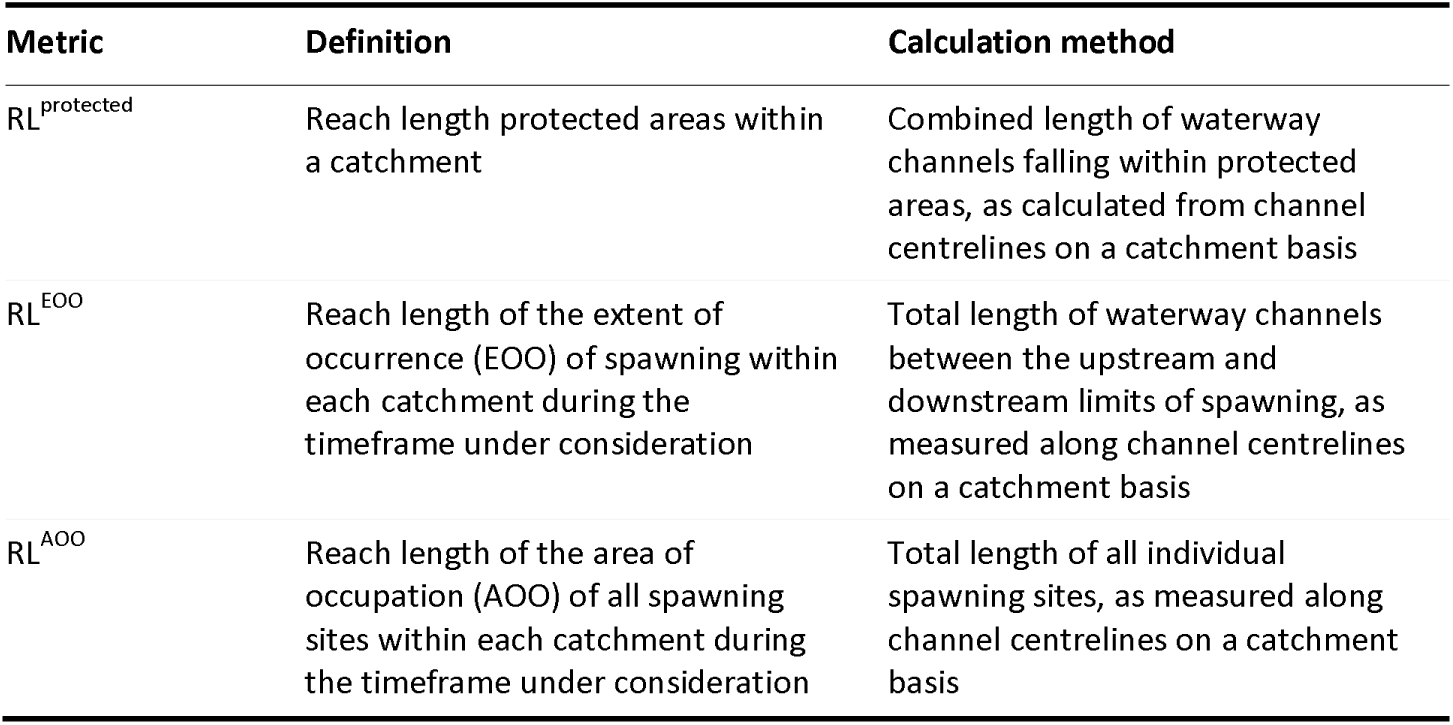
Metrics calculated to evaluate the effectiveness and efficiency of protected area mechanisms for *G. maculatus* spawning habitat.

| Metric | Definition | Calculation method |
| --- | --- | --- |
| RL<sup>protected</sup> | Reach length protected areas within a catchment | Combined length of waterway channels falling within protected areas, as calculated from channel centrelines on a catchment basis |
| RL<sup>EOO</sup> | Reach length of the extent of occurrence (EOO) of spawning within each catchment during the timeframe under consideration | Total length of waterway channels between the upstream and downstream limits of spawning, as measured along channel centrelines on a catchment basis |
| RL<sup>AOO</sup> | Reach length of the area of occupation (AOO) of all spawning sites within each catchment during the timeframe under consideration | Total length of all individual spawning sites, as measured along channel centrelines on a catchment basis |

The effectiveness of each protection mechanism was evaluated as the percentage of post-earthquake RL^AOO^ located within the PA. Efficiency was considered using two ratios: RL^EOO^ to RL^protected^ and RL^AOO^ to RL^protected^. These reflect the size of the area set aside for protection (in terms of reach length) versus the extent of the spawning reach, and the size of the areas actually utilised for spawning respectively. Each calculation was made on a catchment basis at a yearly temporal scale (i.e. 2015 and 2016), and also using the combined data from both years of post-earthquake surveys.

## 3. Results

The three protected area mechanisms provided considerably different RL^protected^ values reflecting their spatial basis (Table 4).. However for each mechanism the RL^protected^ was comparable between catchments. An overlay of each protection mechanism on combined post-quake spawning site data is provided for each of the study catchments in Figure 2.

**Table 4.** Reach length (RL) protected by each of the three protected area mechanisms evaluated in the two study catchments.

| Method | Description of protected area mechanism | Reach length protected (m) |  |
| --- | --- | --- | --- |
|  |  | Avon River | Heathcote River |
| 1 | Network of small protected areas based on known spawning sites | 120 | 80 |
| 2 | Mapped reaches of potential spawning habitat on a catchment basis | 3230 | 3098 |
| 3 | Mapped polygons of predicted spawning habitat coupled with a text description of where in the polygon the protection requirements apply | 19100 | 16600 |

**Figure 2.**
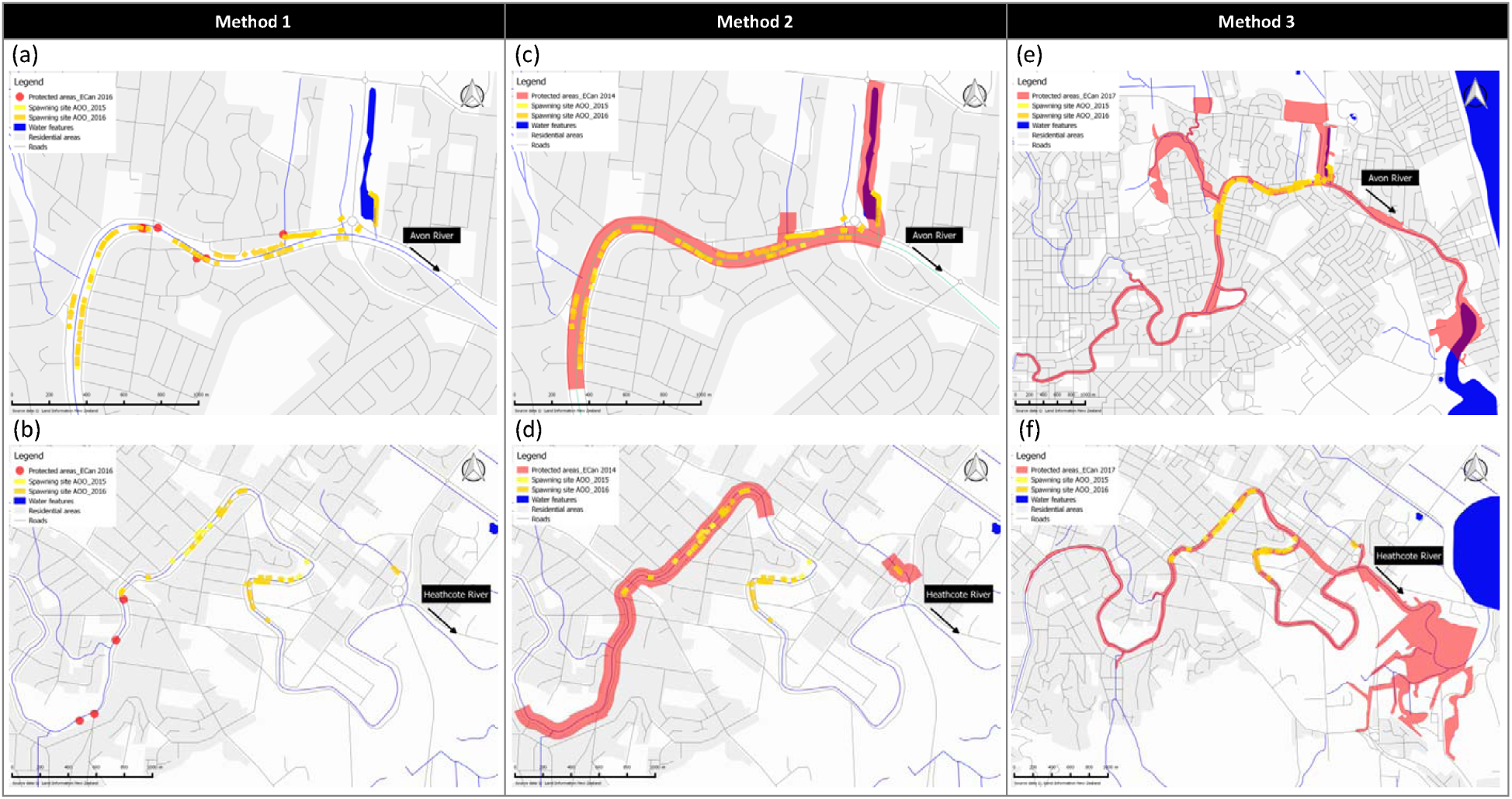
Overlay of the spatial extent of three protection mechanisms found in conservation plans on the footprint of post-earthquake *G. maculatus* spawning sites recorded in 2015 (n = 85) and 2016 (n = 103). (a) Method 1, Avon River, (b) Method 1, Heathcote River, (c) Method 2, Avon River, (d) Method 2, Heathcote River, (e) Method 3, Avon River, (f) Method 3, Heathcote River.

Method 3 was highly effective at protecting spawning habitat, achieving 92.7% protection in the Avon and 100% in the Heathcote using the combined post-quake data (Table 5). The anomoly in the Avon relates to a few spawning sites that occurred outside of the mapped polygon in the vicinity of a small tributary, and this occurred in both years. In the Avon, the effectiveness of method 2 was similar with close to 100% achieved (Table 4). However in the Heathcote, only 69.9% of spawning habitat fell within the protected area and 45.6% in 2016. This reflected the occurrence, in both years, of spawning downstream (Figure 2d). In comparison, the effectiveness of method 1 was low. The percentage of habitat protected ranged from 3.9–14.2% (Table 4). This reflected the extent to which spawning occurred at previously known sites which formed the basis for delineation of the PAs (Figure 2a & 2b).

**Table 5.** Effectiveness of three protected area mechanisms for *G. maculatus* spawning habitat following earthquake-induced landscape change.

| Protection mechanism | Time period | Percentage of habitat protected (% RL<sup>AOO</sup>) |  |
| --- | --- | --- | --- |
|  |  | Avon River | Heathcote River |
| Method 1 | 2015 | 5.4 | 7.5 |
|  | 2016 | 14.2 | 6.3 |
|  | 2015+2016 | 9.3 | 3.9 |
| Method 2 | 2015 | 96.9 | 69.9 |
|  | 2016 | 99.0 | 45.6 |
|  | 2015+2016 | 98.0 | 52.5 |
| Method 3 | 2015 | 96.9 | 100 |
|  | 2016 | 96.5 | 100 |
|  | 2015+2016 | 97.2 | 100 |

In the efficiency evaluation, all of the protection mechanisms were relatively inefficient in terms of land use allocation when the evaluation metric was RL^AOO^ (Figure 3a). For all methods, more than half of the RL^protected^ was allocated to areas that were not utilised for spawning habitat over the study period, even when the areas allocated were very small and targetted at previously known spawning sites. The highest percentage overlap with RL^AOO^ was 47.5% achieved by method 1 in the Avon in 2016. However, when the evaluation metric was RL^EOO^ the percentage overlap results changed considerably. Method 1 achieved a 100% overlap in the Avon in both years but in the Heathcote only 12.5% (Figure 3b). Method 2 achieved 67.6% overlap in the Avon (2016) and 48.7% in the Heathcote (2016), whilst method 3 achieved 11.5% in the Avon (2016) and 17.6% in the Heathcote (2016).

**Figure 3.**
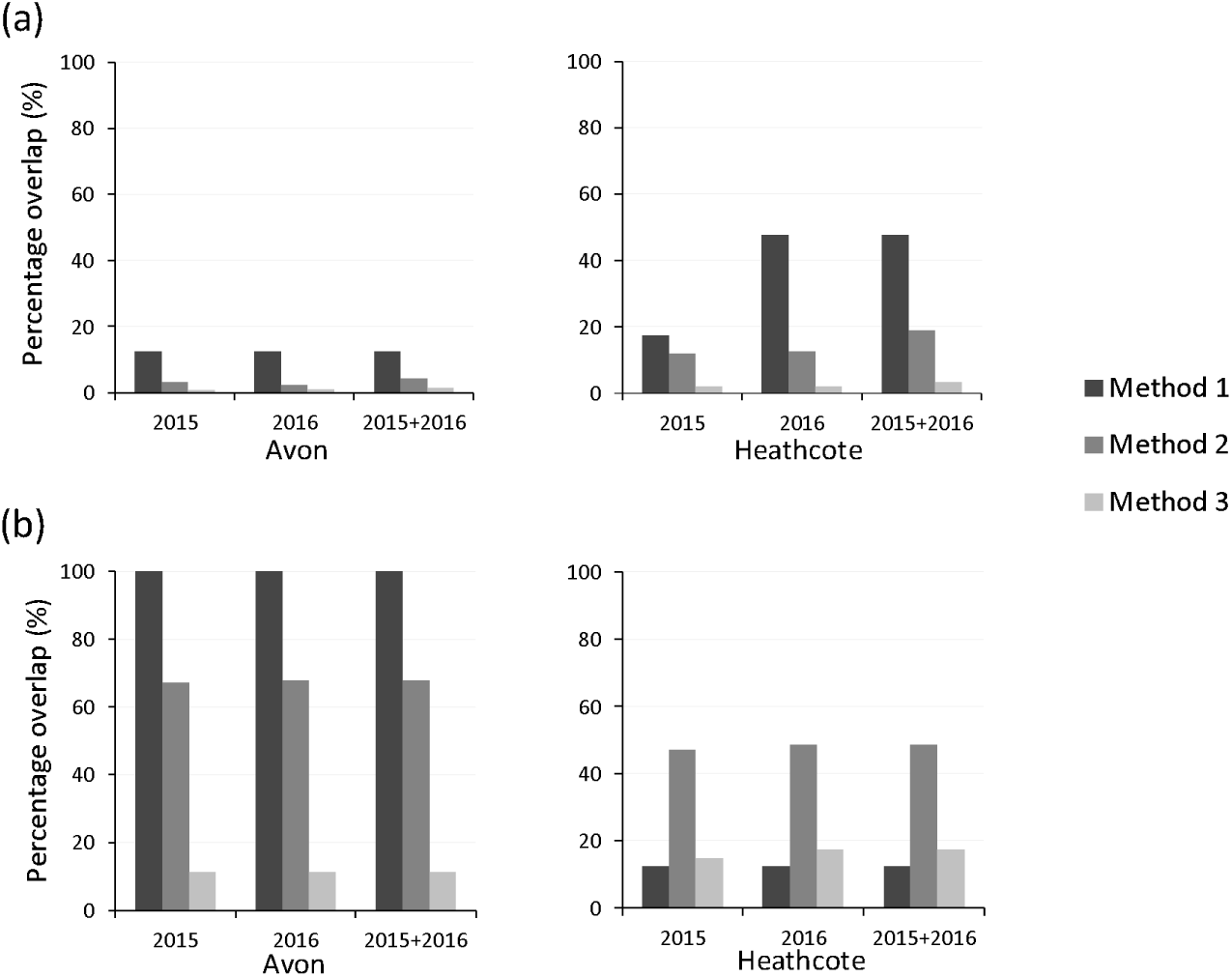
Evaluation of the efficiency of three Category IV protected area mechanisms (Dudley, 2008) in terms of land use allocation using two assessment metrics. (a) percentage of reach length protected (RL^protected^) overlapping the total reach length of areas occupied by spawning sites (RL^AOO^). (b) percentage of RL^protected^ overlapping the reach length of the extent of occurrence of spawning sites (RL^EOO^). In all cases RL is calculated on the centreline of the waterway channel. For each calculation three time periods are considered: 2015, 2016, and combined data from both years.

Comparing these results, method 3 was the least efficient in terms of land use allocation for the purposes of protection in all comparisons in the Avon. However, in the Heathcote method 1 was even less efficient in terms of RL^EOO^ (Figure 3b). This reflected that the protected areas identified were not well located in relation to the areas utilised for spawning (Figure 2). In the Avon, the PAs under the method 1 were much better located with all PAs overlapping the RL^EOO^ (Figure 3b). In terms of RL^AOO^ method 1 also performed better in the Avon versus the Heathcote as a result of the PAs coinciding several of the areas actually utilised. However, even here the efficiency of PA mechanism was rather variable with 47.5% of the RL^protected^ overlapping with spawning sites in 2016 but only 17.5% in 2015 (Figure 3a). This variability is associated with the repeat use of some, but not all, previous used spawning sites between years (Figure 2).

Overall, method 2 produced relatively consistent results in the efficiency comparisions between years. This reflects that the RL^EOO^ was similar in both catchments between years and also located in a similar position in the catchment versus the reaches mapped for protection. Within the RL^EOO^ the total RL^AOO^ was also very similar between years (Avon 386 m^2^ and 410 m^2^, Heathcote 133 m^2^ and 158m^2^ for 2015 and 2016 respectively) despite considerable variation in the location of the sites used each year (Figure 2).

## 4. Discussion

### 4.1 Addressing spatiotemporal variation

Several aspects of *G. maculatus* spawning site ecology are potential sources of spatiotemporal variation. The reported relationship with salinity results in horizontal structuring along the axis of waterway channels in relation to saltwater intrusion (Richardson & Taylor, 2002; Taylor, 2002). This may drive variability in the position of spawning reaches on a catchment scale when coupled with dynamism of river discharges and tidal forcing. Despite that previous studies have highlighted use of the same spawning sites for multiple years (Taylor, 2002), this case was characterised by habitat shift in both catchments in comparison to all known records (Orchard et al., in press). Although the potential effects of salinity changes are not apparent in the literature, this indicates that they may important in relation to perturbations from extreme events or to incremental changes such as sea level rise. However, in relation to this study, a lack of pre-earthquake salinity data for the reaches of interest makes this difficult to confirm directly. The timing of spawning on or soon after the peak of the tide combined with preference for shallow water depths, also leads to vertical structuring of the habitat in relation to water level heights (Benzie, 1968a; Mitchell & Eldon, 1991). Interaction between the waterline and floodplain topography also influences the distance between spawning sites and the alignment of (i.e. perpendicular to) waterway channels. This variation may be considerable where the topography is relatively flat and is a further consideration for effective PA design.

### 4.2 Evaluating PA effectiveness for dynamic habitats

There are at least three aspects of this study that are likely to be applicable to the design and evaluation of Category IV PAs elsewhere. They include the question of PA boundary setting in relation to the habitat to be protected, the need for data to inform this and monitoring strategies to support future evaluations, and practical considerations for identifying boundaries on the ground as required by stakeholders.

Clearly, accuracy is important when setting boundaries for Category IV PAs, yet spatiotemporal variation may hamper acquisition of the necessary data in practice. For *G. maculatus* strong temporal trends are a particular consideration. Variation has been reported in relation to the peak days of activity within a tidal sequence, the tidal sequences preferred in different parts of the country, and months of most spawning activity in the year (Taylor, 2002). International studies have also reported large-scale variation in traits associated with spawning (Barbee et al., 2011). In combination, these aspects suggest that spatiotemporal variability could arise at multiple scales creating practical difficulties for both empirical data collection and model-based approaches for determining habitat distribution. In this case, the study catchments are New Zealand’s best studied spawning areas yet surveys have only been periodic and seldom comprised more than one month in any given year (Taylor, 2002). Consequently, the times of peak spawning activity may not have been captured in the survey record. Identification of the spawning distribution has therefore relied on the compilation of multi-year data despite the potential for confounding factors associated with longer term change.

Albeit that the post-earthquake context represents a major perturbation, the impacts of spatiotemporal variance on PA effectiveness are clearly seen in planning methods 1 and 2. These methods were developed using the planning authority’s up to date information on spawning habitat in both catchments. Particularly in the Heathcote, earthquake-induced habitat shift rendered these methods relatively ineffective. Despite this, regular monitoring and amendment of the same protection mechanism could provide a strategy for maintaining effectiveness and addressing change. However for method 1 the data collection requirements would be onerous to achieve this in practice. This partly reflects reliance on a network of small PAs but also that the detection of spawning sites is difficult (Orchard & Hickford, 2017). The number of PAs identified appears woefully inadequate in light of the post-quake data yet fairly represents results of the monitoring effort that was in place pre-quake. Increasing this to the level of a census-survey for peak spawning months represents a considerably scaling-up of the monitoring programme.

In comparison, method 3 was based on considerably larger PAs and was much more resilient to earthquake changes. In that case, a degree of redundancy was seen as a desirable aspect for resilience (Greer et al., 2015). However, from the perspective of PA evaluation, the three PA mechanisms share similar monitoring requirements. This arises since demonstration of management effectiveness requires information on the values to be protected (Stoll-Kleemann, 2010). Given that monitoring resources are inevitably limited, dynamic environments demand particular attention. In turn this illustrates the widespread need for research on monitoring strategies to inform priorities for data collection and frequency (Teder et al., 2007). Moreover, it exemplifies the need for more management-driven science to close the gap between conservation policy and practice (Knight et al., 2008).

Potential strategies include using abiotic proxies for conservation objectives for which data acquisition is easier thus reducing the burden of repeat measurement (Lawler et al., 2015). Method 3 provides an example of this approach, using a predictive model based on elevation above sea level (Greer et al., 2015). However, the results indicate that its efficiency as a planning method is relatively low since much of the area set aside did not help achieve the stated objectives, and it could not be used as a proxy for outcomes monitoring against the relevant policy objectives. From an ecosystem-based perspective, inefficient planning methods may also hinder potential uses, leading to unnecessary trade-offs (Southworth, Nagendra, & Munroe, 2006). The practical aspects of this relate to the rules that apply within the PA and are designed to confer protection. Where a degree of sustainable use is envisaged within PAs, the specific arrangements for management need to be well matched to intended objectives.

Efficiency may be a particular consideration for Category IV PA evaluation in recognition of the intensity of surrounding resource use that often characterises the management context (Dudley, 2008). In this regard method 2 offered an alternative approach that identified areas of suitable habitat outside of the limits of the known EOO and considered these to be ‘potential’ habitat (Margetts, 2016). These reaches were included in the areas delineated for protection. Essentially this created a buffer around the mapped EOO that served to address limitations in the information available for quantifying known habitat, as well as a providing a degree of redundancy to improve resilience. Although in the Heathcote the post-quake habitat was found to have shifted outside of these areas, they were effective in accommodating the smaller magnitude of change observed in the Avon (Figure 2). Evaluation of method 2 primarily requires information on EOO to determine effectiveness and inform adaptive management. This offers a monitoring strategy that is much less onerous than the census-surveys used in the post-quake studies (Orchard & Hickford, 2017). Method 3 also requires at least this level of monitoring to inform effectiveness evaluation. This suggests that a combination of an evaluation-informed adaptive approach and degree of redundancy could offer an effective and efficient PA strategy for dynamic habitats with regards to land use allocation.

Lastly, this case highlights some practical issues for the visualisation of PA boundaries. In this evaluation, spatial co-occurrence was based on coordinates describing the upstream and downstream extent of spawning sites and polygons describing PAs. In many instances spawning site locations were very close to the PA boundaries as mapped. Unless they were clearly outside of the boundaries, such sites were assessed as being protected with the result being an optimistic view of the extent of the PA mechanism. In reality these boundaries may not be so clear. However, it is important that they *are* clear for the benefit of all stakeholders (Langhammer et al., 2007), and this depends considerably on planning methods. In this case the areas delineated by method 1 were interpreted by stakeholders using a location description and schedule of coordinates (Table 2). This is considered to offer a relatively clear mechanism for implementation of the PA management requirements in practice. Under method 2, the areas for protection were first visualised as lines in Council planning documents (Margetts, 2016) and then subsequently incorporated into ‘Sites of Ecological Significance’ (SESs) in a recent statutory plan (Christchurch City Council, 2015) which is now operative. The visualisation method for plan users is a set of polygons annotated on planning maps appended to the plan (Figure S1a). These SESs have therefore become the PAs of interest and method 2 (as assessed in this study) can be interpreted in relation to *G. maculatus* objectives within these larger areas. However, at the scale of the mapping provided it is difficult to see exactly where the PA boundaries lie in relation to the riparian zone requiring considerable guesswork by plan users (Figure S1b).

Under method 3 the situation is improved by the provision of PA polygons as a public dataset with an online GIS viewer available, in addition to planning maps appended to the relevant plan (Environment Canterbury, 2017). Nonetheless, similar boundary issues arise with regards to the location of the PA in relation to the spatial extent of habitat. The GIS analysis revealed a few spawning sites that were clearly outside of the PA boundary in the Avon, as reflected in effectiveness results of <100% in both years (Table 5), and in general many of the actual spawning locations were again very close to the PA boundary. Furthermore, the habitat may shift a considerable distance from the low flow channel on high water spawning events, and these circumstances are difficult to detect by operators (e.g. management contractors) in the field. Indeed spawning sites were found to have been destroyed by the City Council’s own reserve management contractors subsequent to notification of the relevant statutory plan (Orchard et al., in press). This suggests that better guidance materials, such as interactive maps, may be required to improve PA effectiveness in practice as was recommended in a recent management trial that aimed to avoid such damage to spawning sites (Orchard, 2017). These results also indicate that a buffer should be considered as an aspect of PA design.

### 4.3 Assumptions and limitations

Several assumptions have been made in this evaluation consistent with a focus of the protection of dynamic habitats and the learning available from the unique post-earthquake situation. Most importantly, the focus has been restricted to the spatial basis of protection mechanisms for critical habitat as found in planning documents. In all cases they were assumed to confer protection where spatial overlap occurred. In reality, this also depends considerably on the design of the rules that apply within the PA and aspects such as the provision of compliance monitoring. Also, a conservative approach has been taken in the mapping of PA boundaries and protection assumed to be effective across the whole areas including close the boundaries. In the case of method 2, the width of the riparian zone protected could not be accurately identified and all spawning sites with the protected reach were assumed to be covered. Other limitations of the study include the spatial coverage of post-quake surveys in relation to method 3 since the full extent of those PAs was not directly surveyed. Despite this the spatial coverage of the surveys was extensive in both catchments and the methodology was designed to capturing the upstream and downstream extents of the full habitat distribution (Orchard & Hickford, 2017). Different evaluation results can also be expected in light of new information. In particular the number of spawning events captured in the post-quake survey record is limited. Further spatiotemporal variation may arise from effects such as differing water heights outside of the sampled range, future vegetation change, river engineering impacts, the potential for further ground level changes, and the ongoing influence of sea level rise.

### 4.4 Conclusions

This evaluation was conceived to challenge PA thinking. Firstly, our evaluation extends the discussion of PA management effectiveness towards that of resilience. Although management actions within existing PAs may help increase the resilience of natural resources, the realities of global change create a fundamental challenge that demands a range of approaches (Baron et al., 2009). The PAs involved are small and are best thought of as PA networks under the management of local and regional government entities. Yet in all respects they meet the definition of Category IV PAs and are found nationwide in recognition of their statutory role and origins. Although a focus on critical habitats is just one dimension of protected areas management, it offers a mechanism to help fulfil their potential as management tools through dynamic spatial planning. In particular, attention to relatively fine scales may offer practical opportunities for integrating PA systems into the wider land and seascape (Guarnieri et al., 2016). Small and dynamic PAs have the potential to help fill representation gaps in PA networks as is a critical need in lowland river and floodplain systems (Tockner et al., 2008). Secondly, an understanding of the role of PAs in climate change adaptation processes has been steadily developing but there is much work to be done. For example, new questions to assess the effects of climate change on PAs have only recently been employed in Management Effectiveness Tracking Tool (METT) evaluations despite its long history and widespread use (Stolton & Dudley, 2016). Through investigation of change following an extreme event this study provides insights into similar considerations. Our findings suggest that adaptive networks of well targeted and relatively small PAs could produce an effective mechanism for responding to change thereby contributing to system resilience. Whether new or traditional PAs networks can be adapted along these lines deserves further research. We predict this will become a key topic for environmental planning.

## Acknowledgements

We thank Environment Canterbury and Christchurch City Council staff for information on planning methods and riparian management activities. Funding was provided by the Ngāi Tahu Research Centre and a New Zealand Ministry of Business, Innovation and Employment grant (C01X1002) in conjunction with the National Institute of Water and Atmospheric Research.

---

**Figure S1.**
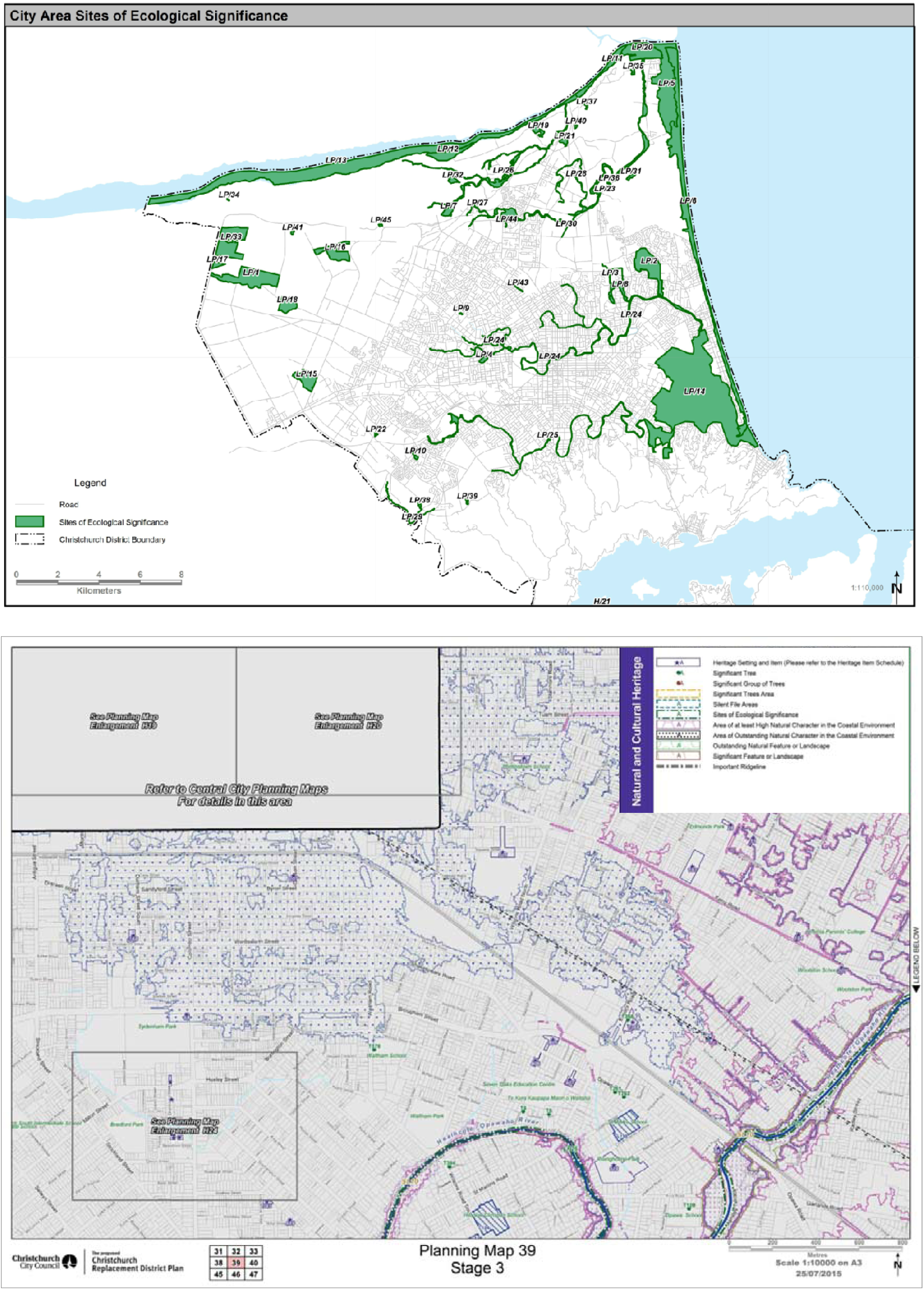
Planning maps showing Sites of Ecological Significance (SESs) in the Christchurch City area (Christchurch City Council, 2015). (a) Schedule Reference Map. (b) Example of detailed planning map. No enlargements are provided for SESs in riparian zones. For brevity only an excerpt of the full legend is shown.

